# Conflict over fertilization underlies the transient evolution of reinforcement

**DOI:** 10.1101/2020.11.10.377481

**Authors:** Catherine A. Rushworth, Alison M. Wardlaw, Jeffrey Ross-Ibarra, Yaniv Brandvain

## Abstract

When two species meet in secondary contact, the production of low fitness hybrids may be prevented by the adaptive evolution of increased prezygotic isolation, a process known as reinforcement. Theoretical challenges to the evolution of reinforcement are generally cast as a coordination problem, i.e. “how can LD between trait and preference be maintained in the face of recombination?” However, the evolution of reinforcement also poses a potential conflict between mates. For example, the opportunity costs to hybridization may differ between the sexes or species. This is particularly likely for reinforcement based on postmating prezygotic (PMPZ) incompatibilities, as the ability to fertilize both conspecific and heterospecific eggs is beneficial to male gametes, but heterospecific mating may incur a cost for female gametes. We develop a population genetic model of interspecific conflict over reinforcement inspired by “gametophytic factors”, which act as PMPZ barriers among *Zea mays* subspecies. We demonstrate that this conflict results in the transient evolution of reinforcement—after females adaptively evolve to reject gametes lacking a signal common in conspecific gametes, this gamete signal adaptively introgresses into the other population. Ultimately the male gamete signal fixes in both species, and isolation returns to pre-reinforcement levels. We interpret geographic patterns of isolation among *Z. mays* subspecies in light of these findings, and suggest when and how this conflict can be resolved. Our results suggest that sexual conflict over fertilization may pose an understudied obstacle to the evolution of reinforcement.

---

> Once the pollen grain has arrived at the stigma, it has made an irreversible move. There should be very intense selection for it to get around whatever barriers the female may erect. — Janzen (1977)

Reproductive interactions present “different evolutionary interests of the two sexes” (Parker and Partridge 1998). This sexual conflict (in the general sense *sensu* Parker 2006) stems from sex differences in the fitness consequences of mating. One sex (often the male) generally benefits from increasing their mating opportunities, while the other sex benefits from choosing mates and/or limiting mating — can result in the evolution of more specific forms of sexual conflict (e.g. harmful mating tactics and mating evasion Rowe and Arnqvist 2005). Because mating and fertilization play a key role in mediating gene flow between divergent populations, sexual conflict can impact the process of speciation. Species boundaries may be strengthened if sexual conflict poses a barrier to gene flow, or weakened if populations evolve mating tactics that can overcome heterospecific reproductive barriers (Parker 1979; Parker and Partridge 1998; Gavrilets and Hayashi 2005; Gavrilets 2014).

The cost of producing low-fitness hybrid offspring can favor the evolution of enhanced reproductive isolation by a process known as reinforcement (Dobzhansky 1937). Reinforcement is generally modeled as the evolution of enhanced prezygotic isolation via female preference and male trait, or trait matching (Kopp *et al*. 2018, although some models have also looked at reinforcement of male mating behavior e.g. (Servedio 2007; Aubier *et al*. 2019)). Such models, generally include trade-off between being attractive to conspecifics and heterospecifics (i.e. species evolve preferences for different trait values), and as such both sexes tend to benefit from assorting with conspecifics and avoiding the production of low fitness hybrids. As such, most reinforcement theory aims to to address the logistical challenge of maintaining a genetic association between trait and preference (Felsenstein 1981), rather than the strategic challenge posed by misaligned interests between the sexes. This is true even in models of polygyny (i.e. when females have multiple mates) because males who match a given female trait/preference are assumed to be less attractive to females with a different trait/preference (Servedio 2006, 2007). However, if the trade-off between conspecific and heterospecific is less severe, missing conspecific mating opportunities can come at a cost for one sex (usually males), while for the other sex (usually females) the cost of producing a low-fitness hybrid often outweighs the marginal benefit of additional matings.

The imbalance between sexes in the opportunity costs of heterospecific mating sets the stage for sexual selection to impact the evolution of reinforcement. For example, Servedio & Bürger (2014) showed that females with preferences misaligned with their species can favor males with traits that do not match the population’s environment, inhibiting the evolution of reinforcement. Similarly, because a male preference results in more competition for mates than does indiscriminate mating (Servedio 2006), reinforcement by male mate choice is more constrained than reinforcement by female choice (Servedio 2007). Likewise, Aubier et al. (2019), found that male preference for conspecifics only evolved if potential male effort toward courting unpreferred females could be reallocated to preferred females. These differences in models of reinforcement by male and female mate choice suggest a conflict in the evolutionary interests of the sexes during the evolution of reinforcement. That is, the benefit of siring low fitness hybrids may exceed the opportunity cost for a male but not for a female; presenting an overall benefit only to males.

The sexual conflict over reinforcement described above is likely particularly severe for reinforcement of postmating prezygotic (PMPZ) isolation, because reproductive effort cannot be reallocated to preferred partners after mating has already occurred (Janzen’s 1977 “irreversible move”, above). From the male perspective, gametes transferred to heterospecific females cannot be redirected, so universally compatible alleles in male gametes will be favored. Whereas from the female perspective, alleles that discriminate against heterospecific gametes in favor of conspecific gametes will be favored.

Motivated by the genetic basis of PMPZ isolation between hybridizing subspecies of *Zea mays* (Mangelsdorf and Jones 1926), we develop a population genetic model to evaluate how this sexual conflict over hybridization can alter the evolution of reinforcement. Our model finds that female requirement of a species-specific signal from male gametes in one “reinforcing” species leads to initial reinforcement of reproductive isolation. But this male signal then adaptively introgresses across species boundaries into the “non-reinforcing species”. Fixation of the male compatibility allele across the metapopulation renders the benefit of discerning female-expressed alleles neutral, resulting in the erosion of reinforcement. Notably, we find a similar outcome when two populations have their own unique incompatibilities, suggesting that this result is attributable to asymmetric costs and benefits experienced by the sexes, and not simply asymmetric cross compatibility.

## Results

### Biological inspiration

Gametophytic factors — pairs of tightly-linked loci expressed in pollen and styles — underlie the PMPZ barrier (Moran Lauter *et al*. 2017; Lu *et al*. 2014, 2019; Wang *et al*. 2018). Counter to the classic reinforcement prediction that reproductive isolation will be highest in areas of sympatry, and reduced in areas of allopatry (Coyne and Orr 1989)), highland teosinte (*Z. m*. subsp. *mexicana*) growing in sympatry with domesticated maize landraces (*Z. m*. subsp. *mays*) shows elevated PMPZ isolation from allopatric maize populations, but no elevation in PMPZ isolation from sympatric maize (Kermicle *et al*. 2006; Kermicle and Evans 2010). Our model is inspired by the function of gametophytic factors, and their puzzling bio-geographic distribution.

Despite this inspiration, our model is neither specific nor fully faithful to the maize/teosinte system. To highlight the generality of our model, we refer to the reinforcing population/species as *reinf*) and the non-reinforcing population/species as *non-reinf*, which roughly represent *Z. m*. subsp. *mexicana* and *Z. m*. subsp. *mays*, respectively. Despite being inspired by a hermaphroditic system, we use the terms “male” and “female” to refer to male and female function, or sperm/pollen and female reproductive tract function, respectively.

### Model overview

We deterministically iterate migration, gamete fusion, and selection, between two populations in secondary contact. See Supp. Text S1 for full description of this iteration, and https://github.com/carushworth/gameto-theory for the R code.

#### Population structure, migration, and pollination/mating

We model two demes (i.e. a two-island model) in two differing selective environments. Every generation, *g*_*non-reinf→reinf*_ of sperm/pollen in the *reinf* environment originates from the *non-reinf* environment. Similarly, *g*_*reinf→non-reinf*_ sperm/pollen in the *non-reinf* environment originates from the *reinf* environment (Figure 1B). Within each environment, sperm/pollen and females meet at random.

**Figure 1.**
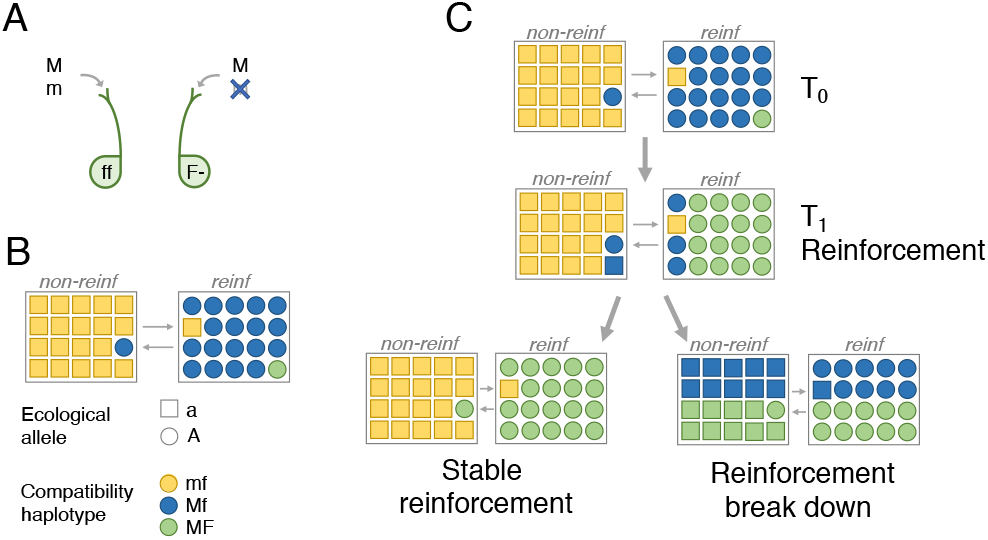
Model dynamics. **(A)** PMPZ incompatibility based on Gametophytic Factors. The dominant *F* allele at the female-expressed locus encodes a fertilization barrier that can only be overcome by the male-expressed compatibility allele *M*. (B) Male gametes disperse between two populations: one reinforcing (*reinf*) and one non-reinforcing (*non-reinf*), each of which face different selective pressures. Colors denote compatability haplotypes (*mf, Mf, MF*), and shapes signify genotypes at a locus underlying divergent ecological adaptation (*a* and *A*). Initially, *non-reinf* is fixed for the compatible *f* female-expressed allele, the incompatible sperm/pollen-expressed *m* allele, and a locally adaptive allele at locus 𝒜 (haplotype *mfa*). *Reinf* is initially fixed for the sperm/pollen compatibility allele *M*, and locally adapted allele *A*, with the female-expressed incompatibility *F* at an initially low frequency. That is, in *reinf* haplotype *MfA* is initially common and *MFA* is initially rare. (C) We run the model from the initial generation *T*_0_ until the *F* allele reaches its equilibrium. If some reinforcement evolves by time *T*_1_ (equivalent to the *F* allele increasing in frequency in *reinf*), two further outcomes are possible: the *M* allele may introgress onto the locally adaptive background and fix in both populations, leading to the breakdown of reinforcement (bottom right panel of **C**); or, the *M* allele may fail to spread in *non-reinf* while *F* continues to spread through *reinf*, completing reinforcement (bottom left panel of **C**).

#### Fertilization

Although mating/pollination within a deme is random, fertilization is controlled by a two-locus PMPZ incompatibility (*sensu* Lorch and Servedio (2007), which is a specific form of a “preference/trait” model, with complete female choosiness (Kopp *et al*. 2018)). The female-expressed locus ℱ is under diploid control. We assume the incompatibility is dominant—i.e. females with one or two *F* alleles discriminate between sperm/pollen alleles, preferentially accepting those with the *M* allele (Figure 1A). Fertilization is random for females homozygous for the compatibility allele, *f*. This notation differs from that in the existing literature on gametophytic factors, which refer to gametophytic factors as haplotypes rather than pairs of genotypes (see Supp. Text S2).

We initially assume no direct fitness cost to either the female incompatibility *F* (e.g. there is no preference cost) or the male compatibility *M*, unless otherwise noted. We further assume that the female incompatibility allele is initially rare in *reinf* (1% frequency) and absent in *non-reinf*, and that the male compatibility allele is fixed in *reinf*, and absent in *non-reinf*, respectively. Variation in the initial frequency of *M* in *reinf*, however, has nearly no effect on the outcome (Fig. S1). Finally, we assume that females expressing the incompatibility genotypes cannot be fertilized by incompatible sperm/pollen. Incomplete penetrance of the barrier results in expected quantitative differences in results, but does not change the qualitative outcomes (Fig. S2).

#### Selection

We model isolation by local adaptation (following Kirkpatrick 2001) with extrinsic isolation driven by *n* local adaptation loci, each noted as 𝒜_*i*_, where the subscript *i* is an arbitrary index of loci. Our primary analyses focus on the case of one locally adaptive locus (i.e. n = 1), an assumption that is relaxed where noted. Selection coefficients are *s*_*non-reinf*_ and *s*_*reinf*_ in environments of the non-reinforcing and reinforcing species, respectively. Fitness *w* is multiplicative within and among loci, *w* = (1 − s_env_)^# maladapted alleles^.

#### Recombination and genome structure

We initially assume a locus order of 𝒜_1_ℳℱ, with recombination rates 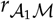 and *r*_ℳℱ_. Local adaptation loci 𝒜_2_ through 𝒜_*n*_ are unlinked to one another and to the ℳ and ℱ loci. After presenting these results, we explore alternative marker orders.

#### A second gametophytic factor

To ensure that our results are not due to asymmetry of variation for female choice in only one population, we then introduce a model with a second unlinked incompatibility locus: 𝒜_*z*_ ℳ_*z*_ ℱ_*z*_. This barrier acts like the first, detailed above, but with initial frequencies in each population reversed. As such, each population is reinforcing and non-reinforcing at a different set of loci.

### Sexual Conflict Leads to Transient Reinforcement

When reinforcement evolves, it is almost always transient. An example of the rise and fall of reinforcement is shown in Figure 2 (Parameter values in legend). Fig. 2A shows that the evolution of substantial reinforcement (Phase 1, Fig. 2A), is ultimately fleeting. Reinforcement begins as the female incompatibility allele, *F*, increases in frequency in *reinf* (Fig. 2B), preventing fertilization by locally maladapted immigrant haplotypes. This maintains both the high frequency of locally adapted (*A*) and male compatible (*M*) alleles in the environment of *non-reinf* (Fig. 2C), and large LD between them (note the large value of 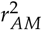 in Fig. 2D,F).

**Figure 2.**
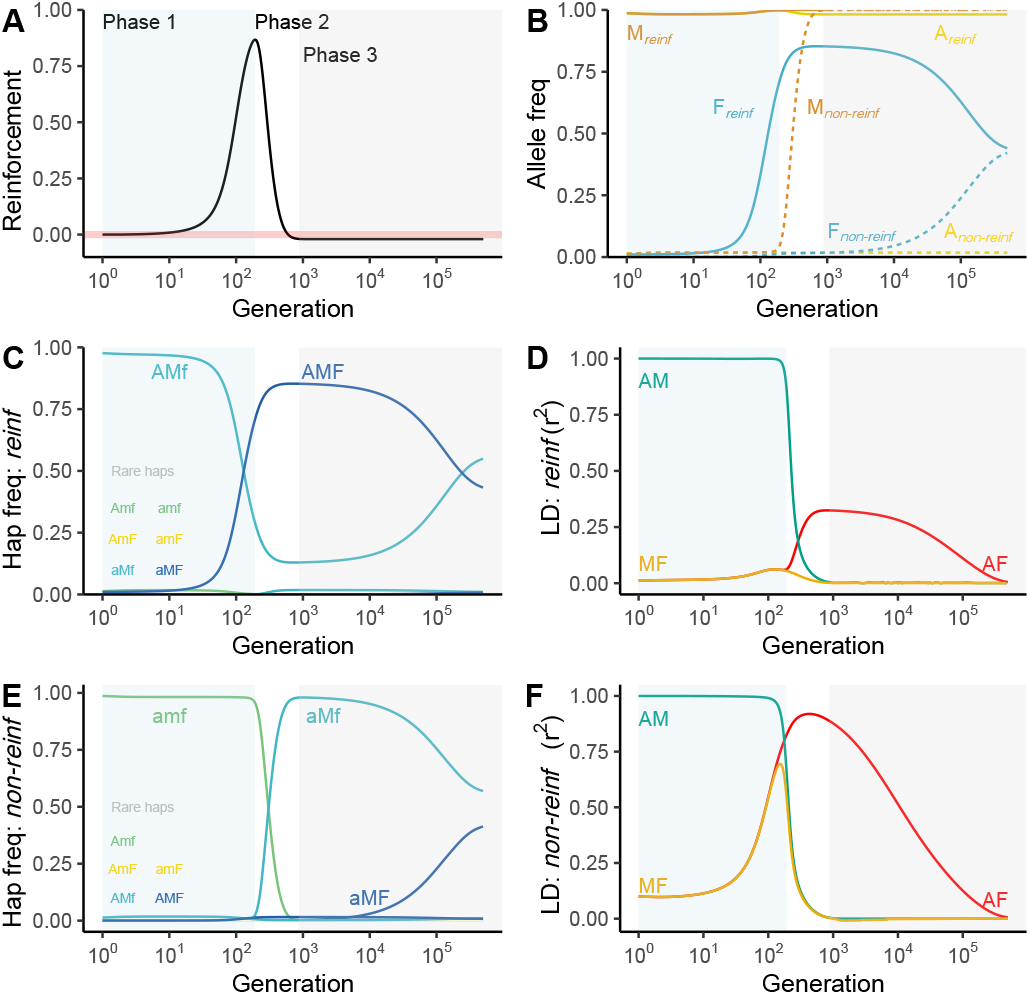
The rise and fall of reinforcement in three phases. A female barrier allele *F* preventing fertilization by *m* gametes spreads in the reinforcing population/species *reinf* (Phase 1: light blue). The compatible sperm/pollen allele, *M*, next introgresses into the non-reinforcing population/species *non-reinf* and spreads (Phase 2: white). After *M* fixes, the barrier *F* slowly disassociates from the reinforcing background, eventually equilibriating in both populations (Phase 3: light grey). **(B)** Reinforcement is transient, building in Phase 1 and breaking down in Phase 2. The pink line shows zero reinforcement. **(B)** Allele frequencies in both populations, with solid and dashed lines showing frequencies in *reinf* and *non-reinf*, respectively. The *F* allele increases in *reinf* followed by the global fixation of *M* and subsequent neutrality of *F*. (**C**) Haplotype frequencies, and (**D**) gametic linkage disequilibrium (LD) between all pairs of loci over time in the environment of *reinf*. (**E**) Haplotype frequencies, and (**F**) Gametic linkage disequilibrium (LD) between all pairs of loci over time in the environment of *non-reinf*. LD is measured as *r*^2^, and all measures describe populations after selection and before recombination. This figure illustrates a single set of parameter values with one adaptive locus. Selection: *s*_reinf_ = *s*_non-reinf_ = 0.75; Migration: *g*_non-reinf→reinf_ = *g*_reinf→non-reinf_ = 0.1; Recombination: *r*_𝒜ℳ_ = *r*_ℳℱ_ = 0.0001; Allele frequencies: *f*_*M*0,reinf_ = 1, *f*_*M*0,non-reinf_ = 0, *f*_*F*0,reinf_ = 0.01, *f*_*F*0,non-reinf_ = 0.

Subsequently, however, the male compatibility allele *M* introgresses into *non-reinf*, eventually recombining onto the *a* background and undermining reinforcement (Phase 2 of Fig. 2). That is, migration of haplotypes from *reinf* into *non-reinf* enables recombination of *M* onto the locally adapted background. As the *aM f* haplotype sweeps through *non-reinf* (Fig. 2E), LD between *M* and *A* decreases in both populations (Fig. 2D,F).

As *M* rises in frequency and eventually fixes across the metapopulation (Fig. 2B), migrant sperm/pollen are no longer rejected, indicated by reinforcement approaching zero in Fig. 2B. At this point, selection against *F* in *non-reinf* weakens until it is completely neutral (see discussion of Fig. 3, below). From then on, *F* slowly equilibriates across populations (Phase 3 of Fig. 2A) as continued migration and recombination between the 𝒜 and ℱ loci decreases LD between them (Fig. 2D,F).

**Figure 3.**
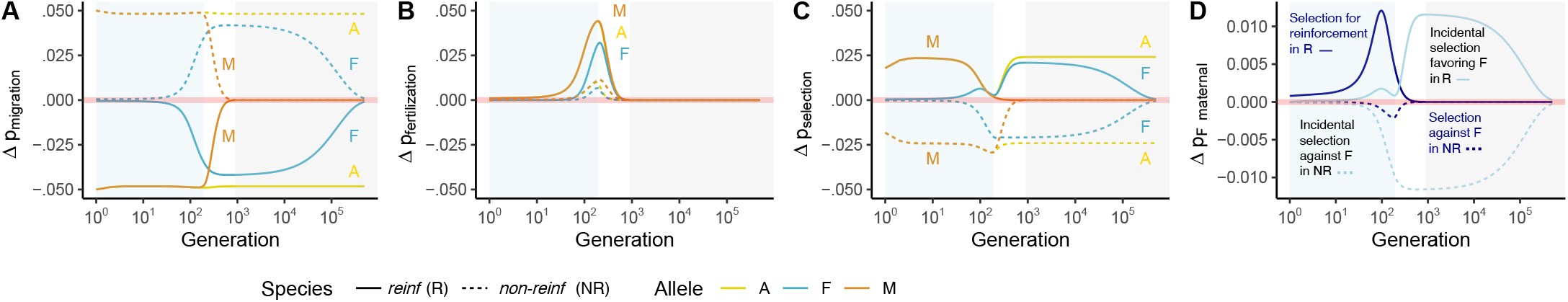
Allele frequency change across the life cycle. Alleles *A, M*, and *F*, which are initially unique to *reinf*, change in frequency across the metapopulation as described in the main text. The transparent thick pink line at zero denotes no change in allele frequency during this life phase. During migration (**A**), alleles native to *reinf* decrease in frequency in *reinf* (solid line, indicated by values below the zero line) and increase in *non-reinf* (dotted line, indicated by values above the zero line). During fertilization (**B**), alleles native to *reinf* (*A, M*, and *F*) increase in both populations. This effect is strongest in the environment of *reinf*, due to a direct fertilization advantage of *M* and the benefit to alleles in LD with *M*. (**C**) Selection in *reinf* and *non-reinf* consistently acts to increase and decrease the frequency of *A*, respectively. Linkage between *A* and *M* results in overlapping lines during phase 1. The transition from phase 1 to phase 2 is marked by a dip in the frequency of *A*, caused by near-fixation of *F* on the *A* background, as migrant haplotypes from *non-reinf* are unable to penetrate *reinf* at *F*’s peak frequency. (**D**) decomposes selection on *F* into two components of allele frequency change. In dark blue we show “selection for reinforcement” (the *F* allele frequency change attributable to preferential fusion with *M*), which enables avoidance of the maladapted *a* allele in *reinf*. In light blue we show the allele frequency change attributable to the incidental gametic phase linkage between *F* and *A*; see Methods for more detail. **Parameter values:** Figures generated with one locally adaptive allele with *s*_reinf_ = *s*_non-reinf_ = 0.75, migration rates, *g*_non-reinf→reinf_ = *g*_reinf→non-reinf_ = 0.1, and recombination rates *r*_𝒜ℳ_ = *r*_ℳℱ_ = 0.0001. Initially, populations are locally adapted (*f*_𝒜,reinf_= 1 − *f*_𝒜,non-reinf_ = 1), with *M* fixed and absent in *reinf* and *non-reinf*, respectively, and *F* at frequency 0.01 in *reinf* and absent in *non-reinf*. Background shading marks phase one (light blue), phase two (white), and phase three (grey) of transient reinforcement, as in Figure 2.

### Allele frequency change across the life cycle

We now show how migration, fertilization and selection drive changes in allele frequencies across the life cycle (Fig. 3). See Supp. Text S3 for exact expressions.

#### Migration homogenizes allele frequencies

The change in allele frequency by migration is the difference in allele frequencies between populations weighted by the migration rate (Eq. S1). This homogenization of allele frequencies (Figure 3A), is seen as the decrease in frequency of all ‘local alleles’ (*A,M*, and *F* always decrease in *reinf* and increase in *non-reinf*).

#### The stylar barrier *F* favors the male compatibility allele *M* and indirectly favors alleles in LD with it

The fertilization advantage of *M* depends on the proportion of incompatible females in the present generation (either heterozygous or homozygous for *F*). For a dominant female incompatibility, this equals 1 − *p* _*f f*_, where *p* _*f f*_, the frequency of females homozygous for the *f* allele, differs from 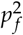 due to non-random fertilization. The increase in frequency of allele *M* (derived in Eqs. S2 and S3) from sperm/pollen to paternally-derived haplotypes equals

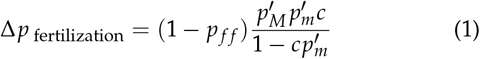

where *c* is the intensity of incompatibility (or choosiness), and the super-script *′* indicates that allele frequencies in sperm/pollen are taken after migration, while female frequencies lack *′* because only sperm/pollen migrate.

In line with this result, Figure 3B shows that in both populations, the male compatibility allele, *M*, increases in frequency during fertilization until it reaches fixation. In addition to directly increasing the frequency of the *M* allele, selection indirectly favors alleles in LD with it (Eq. S4-S5). Because LD among alleles from *reinf* > 0, the *A* and *F* alleles increase in frequency through a fertilization advantage to *M* in both populations (Fig. 3B). This incompatibility system generates a *trans* association between maternally-derived ℱ and paternally-derived ℳ alleles (Eq. S6, and Veller *et al*. 2020).

#### Allele frequency change by natural selection follows standard expectations

Selection increases the frequency of the locally adapted allele at locus 𝒜 in each environment (Fig. 3C, Eq. S7-S8). Likewise, linked selection on *M* and *F* alleles (Fig. 3C) reflects LD with the locally adapted alleles (Fig. 2D and F), with alleles in positive LD with *A* increasing in frequency in *reinf*, and decreasing in *non-reinf* (Eq. S9).

#### Selection favors the female incompatibility in the reinforcing species and disfavors it in the non-reinforcing species

The female incompatibility allele, *F*, which does not itself impact fitness, still deterministically changes in frequency due to its LD with alleles at other loci. This LD is generated by both the causal effect of the allele in mediating non-random fertilization, as well as population structure, historical events, etc. We partitioned the extent to which the increase in frequency of the female isolating barrier *F* is attributable to its causal effect on creating genotypic LD by imposing assortative fertilization (which we call “selection for reinforcement”) versus incidental selection unrelated to the effect of *F* on preferential gamete fusion (see Methods for details).

*F* initially rises in frequency in *reinf* because at the outset it preferentially fuses with sperm/pollen unlikely to contain a locally maladapted allele (Phase 1 Fig. 3D). However, *F*’s persistence once *M* has reached appreciable frequency in *non-reinf* is primarily attributable to linked selection, in that it preferentially exists on locally adapted haplotypes (Fig. 3D).

In *non-reinf, F* is disfavored through both selection against this incompatibility (because *F* is preferentially fertilized by *A*-bearing sperm/pollen) and linked selection acting on the *A* locus (because *F* is in gametic phase LD with the locally maladapted *A* allele). As recombination erodes LD between *M* and *A*, selection weakens against the incompatibility *F* in *non-reinf* due to its effects on non-random fertilization. Indirect selection against *F* in *non-reinf* similarly weakens as recombination erodes LD between *F* and *A* (Fig. 3D).

### Determinants of the strength and duration of reinforcement

We now show how varying parameter values influence the maximum amount (Fig. 4A-D) and duration (Fig. 4E-H) of reinforcement in the face of this conflict.

**Figure 4.**
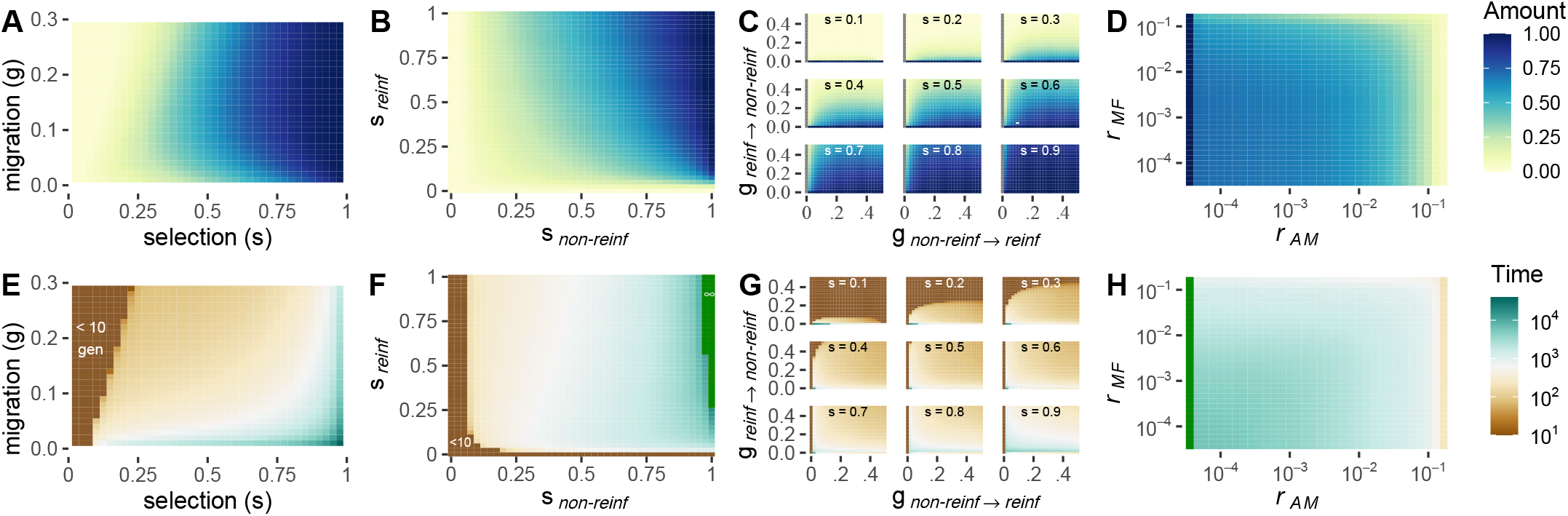
The maximum amount (A – D) and duration (E – H) of reinforcement: Reinforcement as a function of symmetric selection and migration (**A** and **E** with *r*_𝒜ℳ_ = *r*_ℳℱ_ = 10^−4^), different selection coefficients in reinforcing and non reinforcing species’ environments (**B** and **F**, with *g*_non-reinf →reinf_ = *g*_reinf →non-reinf_ = 0.03 and *r*_𝒜ℳ_ = *r*_ℳℱ_ = 10^−4^), asymmetric migration rates across numerous selection coefficients (**C** and **G**, with *r*_𝒜ℳ_ = *r*_ℳℱ_ = 10^−4^), and recombination rates (**D** and **H**, with a symmetric selection coefficient of 0.8 and *g*_non-reinf →reinf_ = *g*_reinf →non-reinf_ = 0.03). Complete or non-transient reinforcement is visible on the far right of figures **F** and **H**, indicated by the darkest green color, and the ∞ symbol in **F**. The amount of reinforcement is quantified as (*p*_[*z*,gen=x]_ − *p*_[*z*,gen=0_])/*p*_[*z*,gen=0]_, where *p*_*z*_ equals the probability of being fertilized by non-migrant sperm/pollen, scaled by the frequency of non-migrant sperm/pollen (“gen” refers to time in generations).

#### Reinforcement often requires strong selection

As selection on the locally adaptive allele intensifies, both the maximum extent (Fig. 4A) and total duration (Fig. 4E) of reinforcement increase. With symmetric selection and symmetric migration, selection on the local adaptation locus must be exceptionally strong for any reinforcement to evolve—e.g. even with *s* = 0.3 only very subtle reinforcement evolves for a very short time. However, other parameter choices—such as asymmetric migration or selection—can mediate the strength of selection required for reinforcement to evolve (see below).

With symmetric migration and asymmetric selection, the strength of selection in *non-reinf* (the population without the stylar incompatibility) generally has a greater effect on the extent and duration of reinforcement than does the strength of selection in *reinf* (Fig. 4B and F, Fig. S3). This is because strong selection in *non-reinf* removes the migrant *MA* haplotype, minimizing opportunities for *M* to recombine onto the locally adapted *a* background.

#### The extent and symmetry of migration mediates reinforcement

With symmetric migration, intermediate migration rates always maximize the extent of reinforcement (Fig. 4A, Fig. S3A), while the duration of reinforcement decreases with the migration rate (Fig. 4E, Fig. S3B), regardless of the selection coefficient.

The effect of asymmetric migration on the extent of reinforcement highlights how migration mediates this sexual conflict. Migration from *non-reinf* to *reinf* favors reinforcement by increasing the number of maladapted immigrants available for heterospecific matings (Fig. 4C). By contrast, increasing migration from *reinf* to *non-reinf* accelerates the introgression of the *M* allele into *non-reinf*, especially at higher migration rates, rapidly undermining reinforcement (Fig. 4G). With unidirectional migration from *non-reinf* to *reinf*, substantial reinforcement can persist for prolonged time periods (Fig. 4C,G).

#### Linkage between female barrier and male (in)compatibility alleles does not strongly impact the amount or duration of reinforcement

Contrary to classic results of reinforcement theory (Felsenstein 1981), linkage between the male and female (in)compatibility alleles, *r*_ℳℱ_, has only a modest effect on the evolution of reinforcement. This result is seen across most selection coefficients and most values of *r*_𝒜ℳ_ (Fig. 4D,H, reproduced in Fig. S5A,D). Reordering the loci in the model does not alter this outcome — that is, the extent and duration of reinforcement is largely insensitive to *r*_ℳℱ_ in models with loci in ℳℱ𝒜 order (Fig. S5B,E).

Instead, linkage between the local adaptation locus, 𝒜, and either ℳ or ℱ loci are critical to the evolution of reinforcement. Marker order, ℳ𝒜ℱ, highlights the impact of recombination between the components of the PMPZ incompatibility complex on both the duration and intensity of reinforcement (Fig. S5C,F). While both *r*_𝒜ℳ_ and *r*_ℱ𝒜_ modulate the level of reinforcement (Fig. S5C), the duration of reinforcement is independent of *r*_ℱ𝒜_ (Fig. S5E), and nearly completely determined by recombination between the male compatibility ℳ and local adaptation locus 𝒜, *r*_𝒜ℳ_. When 𝒜 and ℳ are tightly linked, substantial reinforcement can evolve and last for some time. The strength and duration of reinforcement drops, initially modestly, and then quite precipitously, as the recombination rate increases, with nearly no reinforcement evolving when 𝒜 and ℳ are unlinked (Fig. 4D, H). Selection modulates this effect of recombination (Fig. S4); when selection is very strong (e.g. s > 0.6) some reinforcement can evolve, even when 𝒜 and ℳ are separated by up to a centiMorgan (i.e. *r*_*AM*_ = 0.01).

This result suggests that the rate of recombination between the local adaptation and male compatibility loci, *r*_𝒜ℳ_, underlies the sexual conflict over reinforcement. When *r*_𝒜ℳ_ is high, meaning 𝒜 and ℳ are loosely linked, *M* can more easily recombine onto the locally adapted *a* background, which facilitates its introgression. By escaping from the *A, M* has greater long-term viability in *non-reinf* than it would if it remained associated with this locally maladaptive allele, increasing the male benefit to overcoming the incompatibility.

### The presence of multiple unlinked local adaptation loci allows for (transient) reinforcement

Our results so far suggest that transient reinforcement by PMPZ incompatibilities requires tight linkage between loci underlying incompatibility and a single locus under divergent selection. However, the genetic architecture of local adaptation is often polygenic (Barghi *et al*. 2020).

We therefore investigate if weaker selection at more unlinked loci can allow reinforcement to transiently evolve by setting 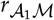 to 0.5, and introducing up to four additional unlinked local adaptation 𝒜 loci. Figure 5 shows that reinforcement can evolve when alternate alleles at numerous unlinked loci experience divergent selection in the two populations. This is consistent with recent work showing that, when numerous loci underlie reproductive isolation, selection on early-generation hybrids acts not against isolated loci, but on phenotypes underlain by pedigree relationships (Veller *et al*. 2019). While the selection coefficients displayed are still quite large, this suggests that weaker selection at many loci could likely result in the transient evolution of reinforcement.

**Figure 5.**
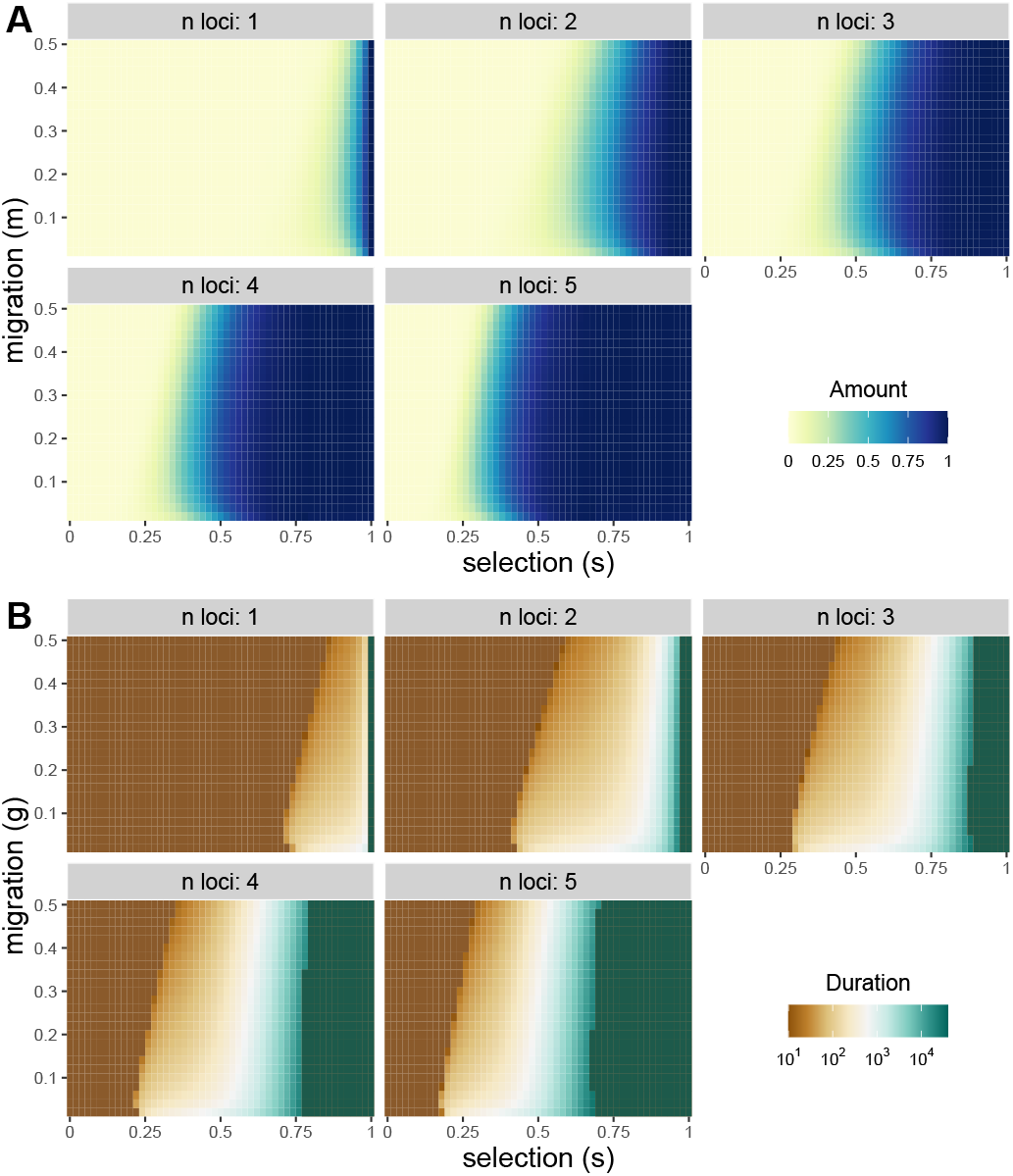
Oligogenic ecological selection: The maximum strength (**A**) and duration (**B**) of reinforcement with ecological selection at *n* loci, where all ecologically selected loci are unlinked to one another and to the gametophytic factor. The selection coefficient *s* against a maladaptive allele is multiplicative within and among loci — e.g. the fitness of an individual homozygous for the locally maladaptive allele at all *n* loci is (1 − *s*)^2*n*^). Migration rate *g* is symmetric and recombination rate *r*_ℳℱ_ = 10^−4^.

### An opposing gametophytic factor does not stabilize reinforcement

To explore the possibility that that the collapse of reinforcement could be prevented by the presence of distinct incompatibilities expressed in each species, we included a model in which both populations are “reinforcing” but by an independent set of loci. Although a second, complimentary, incompatibility allows reinforcement to begin at lower selection intensities and slightly expands the parameter space for which reinforcement stably evolves (Fig. S6A), reinforcement usually remains transient (Fig. S6B).

### A large cost of the male compatibility allele can stabilize reinforcement

We next ask how a cost to the male compatibility allele impacts the evolution of reinforcement; ignoring the question of how such a costly allele could have spread in *reinf*. For illustrative purposes, we limit our focus to exploration in Fig. 2 (also described in legends of Figs. 2 and S8).

A cost, *s*_*M*_, to the *M* can lead to one of three qualitatively different outcomes. If its cost is sufficiently small (*s*_*M*_ < 0.01 in our example) it rapidly and adaptively introgresses and undermines reinforcement as shown above. A larger cost to *M* (0.01 < *s*_*M*_ < 0.08) results in an equilibrium level of reinforcement (i.e. partial reproductive isolation (Servedio and Hermisson 2020)) wherein the cost of *M* is balanced by a benefit of heterospecific fertilization to *M*-bearing pollen/sperm. An even larger cost to *M* (*s*_*M*_ > 0.08 in our example) prevents the introgression of this allele altogether, resulting in stable reinforcement. In sum, a sufficiently large cost to the male compatibility allele can stabilize the evolution of reinforcement, but a small cost does not.

## Discussion

For decades, researchers have presented theoretical and empirical challenges to the process of reinforcement, starting with a foundational paper by Felsenstein (1981) that identified recombination as a critical challenge to reinforcement. Since then, a large body of theory has investigated the circumstances that permit, or hinder, the evolution of reinforcement (reviewed in Kirkpatrick and Ravigné 2002). Despite its potential role in hampering speciation (Parker and Partridge 1998), however, sexual conflict over hybridization has received relatively little attention in the literature. When it is mentioned, sexual conflict over hybridization is included only as a brief aside in papers concerning the role of sexual conflict in speciation more broadly (Parker and Partridge 1998; Gavrilets and Hayashi 2005; Gavrilets 2014).

Here, we identify the transient dynamics generated by sexual conflict over reinforcement and the evolutionary traces it leaves behind — namely the adaptive spread of female barriers in one species/population and the adaptive introgression of male compatibility alleles into the other. These results provide a rich set of predictions and interpretations of empirical patterns that were absent from previous game theoretic (Parker and Partridge 1998; Gavrilets and Hayashi 2005) and verbal (Coyne and Orr 2004) models.

In our model, sexual selection favors sperm/pollen traits that overcome a heterospecific female barrier. This poses a conflict: females are selected to avoid the production of maladapted hybrids, while sperm (or pollen) that increase their fertilization success will generally be favored. The breakdown of reproductive isolation is marked by the rapid adaptive introgression of the male compatibility trait into *non-reinf*, following recombination onto the locally adapted haplotype. Backmigration of this allele into *reinf* hastens its fixation across populations. The final step in the model is the slow homogenization of the female barrier allele across demes, as the male compatibility allele fixes in both, rendering the female choice allele ineffective. Ultimately, we show that barriers acting at different stages of hybridization can affect how reinforcement proceeds. Below we discuss the relationship of our results to previous theory, implications for the process of reinforcement, and empirical implications for hybridizing taxa, including *Zea mays* subspecies.

### Theoretical context and predictions for reinforcement

Previous models of reinforcement treated the sexes interchangeably (Felsenstein 1981), or assumed assortative mating under female control (Lande 1981; Servedio and Kirkpatrick 1997; Kelly and Noor 1996), (but see Spencer *et al*. 1986; Servedio 2007; Aubier *et al*. 2019 for models with male choice), either by “matching” or “preference/trait” mechanisms of assortative mating (Kopp *et al*. 2018). Both matching and preference/trait models induce a trade-off between heterospecific and conspecific mating: a male with a trait favored by heterospecific females will have limited mating success with conspecific females.

While numerous studies have addressed the role of introgression in reinforcement (e.g. Sanderson 1989; Liou and Price 1994; Kirkpatrick and Servedio 1999; Servedio 2000; Matute 2010a), a distinguishing feature of our model is the mechanism of non-random fertilization, which we model as a PMPZ incompatibility functioning as a gametophytic factor in *Zea mays* (Kermicle 2006) and PMPZ barriers in broadcast spawners (e.g. Lessios 2007). In this special class of preference/trait model, introgression of a male compatibility allele is facilitated by the fact that it does not prevent heterospecific mating—that is, by definition our model lacks a trade-off between con- and heterospecific mating (i.e. no reallocation *sensu* Aubier *et al*. 2019). As such, (in)compatibility-type mating interactions can result in the transient evolution of reinforcement, while other matching or preference/trait models cannot.

### Implications for reinforcement by pre-vs post-mating barriers

Our model assumes that being attractive to one species does not come at a cost of being attractive to ones’ own species. How does this assumption likely map onto to cases of reinforcement by pre- and postmating isolation mechanisms?

#### Postmating prezygotic isolation

In our model, sperm/pollen compatibility with heterospecific females does not come at a cost to successful mating with conspecifics. To the extent that PMPZ barriers function similarly to those in our study, our model suggests the transient reinforcement by PMPZ barriers. However, physical and/or biochemical properties of PMPZ interactions may minimize the opportunity for interspecific sexual conflict by enforcing a trade-off between overcoming a heterospecific barrier and successfully fertilizing conspecifics. This would allow for the evolutionary stability of reinforcement, consistent with our results from our model that showed that a sufficiently large cost to pollen/sperm compatibility could stabilize reinforcement. For example, as Howard (1993) argued, and Lorch & Servedio (2007) showed, a preference for conspecific sperm can stably evolve to minimize heterospecific fertilization. Thus conspecific sperm precedence in competitive fertilization likely allows stable reinforcement by PMPZ isolation (as shown by e.g. Castillo and Moyle 2019). Likewise, mechanistic features of non-competitive fertilization can also induce a trade-off between inter- and intraspecific crossing success. For example, if pollen must grow an optimal distance to fertilize an ovule (as observed in interspecific *Nicotiana* crosses Lee *et al*. 2008), success on both hetero- and conspecific styles is impossible.

#### Premating isolation

In principle, our model could apply to the reinforcement of pre-mating isolation, as well as postmating isolation, so long as there is no trade-off in attractiveness to each species. The are numerous examples of premating traits that may increase heterospecific reproductive success without trading off conspecific attractiveness. For example, divergence in male competitive ability between two lineages of the common wall lizard (*Podarcis muralis*) results in asymmetric hybridization (While *et al*. 2015; MacGregor *et al*. 2017). Likewise, loci underlying plumage traits appear to be asymmetrical and adaptively introgress from one to another subspecies of the red-backed fairy-wren, *Malurus melanocephalus* (Baldassarre *et al*. 2014), presumably because this trait confers an advantage in extra-pair copulation due to sensory bias (Baldassarre and Webster 2013). In a classic example of this phenomenon, plumage of the golden-collared manakin *Manacus vitellinus* appears to adaptively introgress into *Manacus vitellinus* upon secondary contact (Parsons *et al*. 1993), and this spread is likely mediated by female choice (Stein and Uy 2006). However, it does not appear that the female preference in any of these cases initially arose as a mechanism of reinforcement.

### Sexual conflict and sexual selection undermines reinforcement

Our model shows that the common phenomenon of sexual conflict (see Arnqvist and Rowe 2005, for examples), wherein male and female interests are misaligned, can erode reinforcement by PMPZ incompatibilities. This role for sexual conflict in removing species boundaries runs counter to the conventional role it is thought to play in speciation (Van Doorn *et al*. 2001; Gavrilets and Waxman 2002; Rice 1996, but see Parker and Partridge (1998)). Previous theory (Gavrilets and Waxman 2002) and experiments (Palopoli *et al*. 2015), as well as natural patterns of reproductive isolation (Brandvain and Haig 2005; Ting *et al*. 2014) and diversification rates (Arnqvist *et al*. 2000) suggest that independent coevolutionary arms races between male and female traits in two incipient species can pleiotropically result in behavioral or mechanical isolation. In this manner, intraspecific sexual conflict was thought to be an “engine of speciation” (Rice *et al*. 2005). By contrast, we show that interspecific conflict between the sexes over fertilization hampers speciation. This highlights an under-appreciated challenge to reinforcement by PMPZ barriers. Broadly, our results align with studies suggesting that incompatibilities (Bank *et al*. 2012), especially those with a transmission advantage (Meiklejohn *et al*. 2018), can adaptively introgress across species boundaries.

Servedio & Bürger (2014) found that Fisherian sexual selection can undermine reinforcement. Specifically they found that migration of female preference alleles provide a mating advantage to otherwise locally maladaptive heterospecific male traits by sexual selection, undermining reinforcement. In contrast, our model shows that the benefit of siring low fitness hybrids, when the alternative is missed fertilization opportunities that result in no offspring, can also undermine the evolution of reinforcement. Counter to Servedio & Bürger, we find that the female incompatability remains at low frequency in the non-reinforcing species for a long time. As such, our models make contrasting predictions: Servedio & Bürgers model predicts that alternative female preferences will coexist in both populations early in the evolutionary process, while our model predicts that female preference for the “wrong” species signal will only become common in both populations very late in the process of genetic exchange.

Our model can be seen as a specific instance of the lek paradox (Borgia 1979), as female preference ultimately erodes variation for a male trait. Following previously proposed resolutions of the lek paradox (e.g. Rowe and Houle 1996), Proulx 2001 suggested that a female preference for an indicator of paternal fitness (e.g. sperm competitiveness) could act as a “single allele” mechanism of reinforcement (sensu Felsenstein 1981). Under a model of local adaptation, however, the efficacy of this resolution depends on the details of population structure. In our model, rather than being resolved, the lek paradox leaves a signature of the “ghost of reinforcement past.”

### Empirical implications, predictions, and interpretation of current observations

Our model, based on a well-characterized PMPZ incompatibility, shows that reinforcement by such mechanisms is precarious. As such, to the extent that such barriers do not incur a tradeoff between conspecific and heterospecific fertilization success, we predict that reinforcement by PMPZ barriers should be rare. Thus, the finding that gametic isolation in broadcast spawners is not the product of reinforcement (contrary to initial claims e.g. (Geyer and Palumbi 2003), discussed in Vacquier and Swanson 2011), as well as meta-analyses showing that PMPZ isolation does not differ between sympatric and allopatric species pairs in *Drosophila* (Turissini and Matute 2017) or across three angiosperm genera (Moyle *et al*. 2004), are consistent with our model. Still, negative evidence is not necessarily evidence for the negative.

As such, the few documented cases in which PMPZ barriers are reinforced allow for better evaluation of its predictions. Specifically, we predict that reinforcement by PMPZ barriers should often involve certain characteristics, which are consistent with the empirical literature. These include: *(1)* recent sympatry, so that the male barrier has not yet increased in frequency (e.g. Poikela *et al*. 2019); *(2)* a trade-off between male success in overcoming inter- and intraspecific postmating barriers, as is found in preference/trait mechanisms (e.g. Castillo and Moyle 2019); *(3)* unidirectional gene flow (e.g. Kay 2006);, and/or *(4)* exceptionally strong postzygotic isolation, such that gene flow is very rare, as seen in *D. yakuba* and *D. santomea* (e.g. Turissini and Matute 2017). However, that reinforcement by PMPZ isolation in *D. yakuba* and *D. santomea* is difficult to reconcile with our model, as the pair have a stable hybrid zone, no evidence of conspecific sperm precedence, and bidirectional hybridization (Matute 2010b). Nonetheless, our model suggests a plausible evolutionary mechanism for existing cases of reinforcement by PMPZ isolation, and generates specific hypotheses to be tested.

### Predictions for maize and teosinte

*Zea mays* ssp. *mays* and *Zea mays* ssp. *mexicana* grow in close sympatry and hybridize (Hufford *et al*. 2013). Evidence for genome-wide selection against admixture despite adaptive introgression of some teosinte loci into maize (Calfee *et al*. 2021), is consistent with the idea that hybrids are often disfavored — perhaps because hybrids are removed from maize fields by anthropogenic weeding, and maize traits expressed in teosinte environments are likely maladaptive (Wilkes 1977; Hufford *et al*. 2012, although clear cases of adaptive introgression and deliberate hybridization by farmers exist).

This system has all the ingredients necessary for reinforcement — the occurrence of gene flow, the presence of a stylar incompatibility in teosinte sympatric with maize, and the reduced but non-zero fitness of hybrids. However, the elevated elevated pollen discrimination exhibited by highland teosinte sympatric with maize (Moran Lauter *et al*. 2017; Lu *et al*. 2014, 2019; Wang *et al*. 2018; Lu *et al*. 2020) is surprisingly ineffective in preventing fertilization by sympatric maize landraces (Kermicle *et al*. 2006; Kermicle and Evans 2010), against whom selection for reinforcement should be strongest.

Our model explains this observation as the initial evolution of reinforcement (i.e. a stylar barrier sweeps through teosinte) followed by the adaptive introgression of teosinte pollen compatibility alleles into maize. Notably, alternative explanations for this pattern are insufficient. For example, this pattern is not simply attributable to the loss of isolation upon secondary contact, because allopatric teosinte do not reject maize pollen (Kermicle *et al*. 2006; Kermicle and Evans 2010). Nor can this be explained by complex speciation, in which teosinte sympatric with maize would be more recently diverged from maize than are allopatric teosinte, as this is incompatible with both genetic evidence and the history of maize domestication (Ross-Ibarra *et al*. 2009). We suggest that in most sympatric populations, at most gametophytic factors, the stylar fertilization barrier (the *F* allele) rapidly swept through teosinte (Phase 1 in Figure 2), and the pollen compatibility allele (*M*) adaptively introgressed into sympatric highland maize landraces (Phase 2 in Figure 2).

### Caveats

Our model made many simplifications and abstractions for tractability and generality. Most notably, we assumed only two populations, a single gametophytic factor, and a simple multiplicative fitness function across a small number of divergently selected loci. Our results show that the architecture of adaptive differentiation, and the linkage between locally adaptive alleles and PMPZ incompatibilities modulate the rise and fall of reinforcement. Across taxa, adaptive differentiation can be controlled by few (Selby and Willis 2018; Lowry and Willis 2010) or many (Yeaman 2013) loci, and linkage between locally adaptive alleles and PMPZ incompatibilities is biologically variable, and rarely known. In maize, evidence is mixed — one gametophytic factor *tga1* is tightly linked to a domestication locus *su1* (Wang *et al*. 2005; Whitt *et al*. 2002) which likely experiences divergent selection. However, the other two known gametophytic factors are far from loci under strong divergent selection.

We further assume that male compatibility allele is initially common and stylar incompatibility is initially rare in *reinf*, and we do not address the origins of gametophytic factors. While the initial divergence of these allele is outside the scope of our model, it could be explained by pleiotropy, selection to prevent polyspermy (a known risk to embryo viability in maize; Grossniklaus 2017), or Fisherian runaway selection (as proposed by Jain 1967, for gametophytic factors). Pleiotropy may explain the initial evolution of gametophytic factors in *Z. mays*, as they are members of the multi-function pectin methylesterase (PME) and pectin methylesterase inhibitor (PMEI) gene families (Moran Lauter *et al*. 2017; Lu *et al*. 2014, 2019, 2020) and could be favored by mechanisms related to other functions. Notably, a subclass of PMEs contain both PME and PMEI domains, providing a potential explanation for tight linkage of the *M* and *F* alleles (Tian *et al*. 2006).

## Conclusions

We find that considering the role of sexual conflict — a mismatch between optimal fertilization outcomes of each sex — in reinforcement generates novel predictions and may explain numerous patterns in nature. Our results are particularly relevant to potential cases of reinforcement by gamete recognition in plants, as well as broadcast spawners (e.g. Lysin/VERL in abalone (Swanson and Vacquier 1998) or Bindin/EBR1 (Metz *et al*. 1994) in sea urchins), and even cases of internal fertilization in which premating isolation is inefficient, costly, or otherwise unpreferred (Turissini *et al*. 2017). In these situations, we predict that reinforcement by PMPZ will be rare, transient, or involve a trade-off between heterospecific and conspecific fertilization (i.e. some mechanism of reallocation). Finally, although our model is developed specifically for interactions between haploid male gametes and diploid females, similar dynamics could arise for premating barriers with a similar genetic architecture and lacking reallocation of male reproductive effort.

## Materials and Methods

### Quantifying reinforcement and its duration

We summarized our results by quantifying the duration and maximum extent of reinforcement. We quantified the amount of reinforcement at generation *g* as (*p*_[*z*,gen=g]_ − *p*_[*z*,gen=0_])/*p*_[*z*,gen=0]_. *p*_*z*_ equals the probability of being fertilized by non-migrant sperm/pollen, scaled by the frequency of non-migrant sperm/pollen. We quantified the duration of reinforcement as the number of generations for which the amount of reinforcement was greater than 0.05.

### Computer Code

All code is written in R (R Core Team 2020) and is available at https://github.com/carushworth/gametotheory. We generated figures with the ggplot2 (Wickham 2016) and cowplot (Wilke 2020) packages, and used the dplyr (Wickham *et al*. 2020) package to process numeric results.

### Partitioning selection

All selection for or against the female incompatibility allele, *F*, is indirect, as it does not itself impact fitness. That is, selection impacts the frequency of an allele at the ℱ locus, not because of its effect on fitness, but because of its genetic background (i.e. linkage disequilibrium between *F* and *A*). Each generation, some of the LD between *F* and *A* is immediately attributable to either (a) population structure, historical events, etc. (primarily by cis-LD) or (b) to the causal effect of the *F* allele in generating genetic associations by the gametes permitted fertilization (primarily via trans-LD; see subsection *The generation of trans linkage disequilibrium during fertilization* in the Mathematical Appendix S3). See Veller (2020) for a discussion of how LD in *cis* and *trans* contribute to Fisherian sexual selection.

Thus the change in frequency in *F* is attributable to both its circumstance (which we call “incidental selection”) and its causal effect on generating a non-random association (which we call “selection for reinforcement”). We aim to partition total selection for (or against) the *F* allele into incidental selection (unrelated to the effect of *F* on non-random fertilization, Δ*p*_*F*,Linked_) and selection for reinforcement (the causal effect of *F* on its selective trajectory, Δ*p*_*F*,Reinforcement_). In this exercise, we ignore the change in frequency of paternally-derived genotypes (which includes migration, fertilization, and selection), as none of this change is plausibly attributable to the *F* allele. We include the subscript *mat* with each variable to remind readers we are focused on the maternally-derived *F* alleles.

We first compute the difference in allele frequency between maternally-derived haplotypes in offspring after vs. before selection as Δ*p*_*F*,mat_ directly from our results. Δ*p*_*F*,mat_ = *p*_*F*,mat-derived after sel_ − *p*_*F*,mom_. We then decompose Δ*p*_*F*,mat_ into components of reinforcing and linked selection:

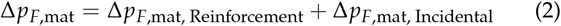

Each generation we find Δ*p*_*F*,mat, Incidental_ by calculating Δ*p*_*F*,mat_ under the counterfactual case of random fertilization. We then find the change in frequency of *F* by selection for reinforcement Δ*p*_*F*,mat, Reinforcement_ by rearranging Eq. 2.

## Acknowledgments

This work was funded by NSF award #1753632 to J Ross-Ibarra, MMS Evans, and Y Brandvain. We are grateful to Robin Hopkins and Maria Servedio, whose comments greatly improved the manuscript, and to Jerry Kermicle for helpful conversation. We would like to acknowledge Felix Andrews for the smorgasbord of statistical advice, although we did not follow it.

## Supplement

### Supp. Text S1: Pseudo-code for our model

We forward iterated genotype frequencies over time, not by iterating a numerical equation, but rather by deterministic programmatic iteration of a the evolutionary process. R scripts are included as a supplemental file, but here we provide pseudo-code to describe the process to the reader.

#### 1. Initiation

Beginning with user-specified initial allele frequencies in each population, as well as additional parameters (e.g. the number of unlinked, local adaptation loci), we build initial genotype frequencies, assuming linkage equilibrium and random mating within each population.

#### 2. Iteration

Every generation, we iterate the evolutionary process, for both populations — one population at a time — by

a. *Meiosis in males:* Generating haploid sperm/pollen grans from diploid genotypes by meiosis, using genotype frequencies in the previous generation.
b. *Migration:* Making a sperm/pollen pool with haplotype frequencies 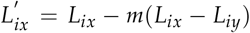. Recall that *m* denotes the proportion of sperm/pollen in population *x* which migrated from population *y*.
c. *Pollination and Fertilization:* We assume that, after migration, sperm/pollen from the falls randomly on styles (or female gametes) within the population, so the frequency of mating between a given sperm/pollen haplotype and female genotype (with frequencies from this population after the conclusion of the previous generation) is their product. Fertilization is non-random, with *c* indicating the strength of discrimination against incompatible *m* alleles. For a given diploid female genotype, the frequency of paternal genomes with alleles *A* and *a* are *p*_*A*_ and *p*_*a*_ × (1 − *c*), respectively, with each divided by the proportion of sperm/pollen compatible with that female genotype (all pollen for *f f* genotypes and (1 − *c*) × *p*_*M*pollen after migration_ for *F f* and *FF* genotypes).
d. *Female meiosis and syngamy:* Finally, females undergo meiosis and zygotes are generated at random, conditional on compatibility between female/stylar genotypes and sperm/pollen haplotype (above).
e. *Selection:* We count the number of alleles at all A loci that mismatch the environment, *n*_maladapt_, and calculate the fitness of each genotype as (1 − *n*_maladapt_)^*s*^. We calculate mean fitness as the product of genotype fitness and genotype frequency, summed over all genotypes. Finally, we find the frequency of each genotype after selection as the product of its frequency before selection and its fitness, divided by mean fitness.
f. *Return summary:* We return a vector of genotype frequencies, as well as the extent of reinforcement in this population in this generation (as described in the Methods). Optional returns are available (e.g. partitioning allele frequency change across the life cycle, and evaluating “counterfactual scenarios” for selection for reinforcement and linked selection on *F*), which we only used for a few exemplary scenarios to minimize computational and storage expense.
g. *Repeat or conclude:* After conducting the steps above for both populations, we store the new genotypes’ frequencies, as well as other quantities of interest. The simulation then returns to (a) if stopping criteria have not been met, or concludes if so. By default, stopping criteria are: simulation length of at least 1000 generations and genotype frequencies at equilibrium (identified when the sum of the absolute value of genotype frequency change across all phased genotypes in both populations is less than 0.00001).

#### 3. Conclusion

At the conclusion of the simulation, we return phased 𝒜ℳℱ genotype frequencies after selection in each population for every generation, the extent of reinforcement every generation, as well as optional summaries mentioned above (see code for all options).

### Supp. Text S2: Notation for gametophytic factors

Our notation throughout differs from the existing literature on gametophytic factors. In this literature, each of the three known gametophytic factors (Ga1, Ga2, and Tcb1) is represented as a single locus with three haplotypes (e.g. Moran Lauter *et al*. 2017). The “strong”‘ haplotype, indicated by *s* or *S*, contains both functional alleles from this study (*F* and *M*). The haplotype with only the pollen-expressed compatibility allele is referred to as *m*, and is equivalent to the *Mf* genotype in the present study. The non-functional allele is referred to in the literature in lowercase (e.g. ga1, ga2, tcb1), and is equivalent to the *mf* genotype in our study. The *mF* genotype is not known in the wild.

### Supp. Text S3: Mathematical Appendix

We present standard recursion equations to provide some intuition for our results, but note that we did not use these recursions to generate our figures, which come from a iteration of the process rather than an explicit recursion of 27 unphased (64 phased) genotype frequencies in two non-randomly mating populations. Nor do we provide quasi-linkage equilibrium or Hardy-Weinberg equilibrium approximations as our focus is on the exact results provided in the main text. As such, the equations below, which largely recapitulate standard results from population genetic theory, are meant to provide some mathematical intuition for our results.

#### Migration

All alleles change in frequency by migration in the same manner. Allele frequencies in pollen at the *i*^*th*^ locus in the focal population *x* after migration, 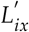, equal

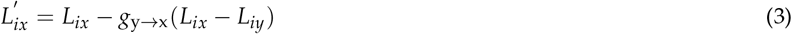

Where the prime ′ denotes the frequency in sperm/pollen after sperm/pollen migration and *L*_*iy*_ is the allele frequency in the other population. From Eq 3, the change in allele frequency at the *i*^*th*^ locus in sperm/pollen grains by sperm/pollen migration equals 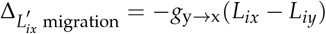, a standard result in population genetics.

#### Fertilization

We assume both no sperm/pollen limitation, and an infinite supply of sperm/pollen, such that the frequency of a sperm/pollen haplotype on a female/style equals the frequency of this sperm/pollen haplotype in the local environment. Under this model, selection during fertilization changes the frequency of alleles in paternally-derived haplotypes relative to their frequency in sperm/pollen after migration, and does not impact allele frequencies of maternally-derived haplotypes.

After mating, the frequency of paternally-derived haplotypes bearing the sperm/pollen allele is *p*_*M*″_, with the double prime denoting that frequencies are calculated after migration and fertilization. This value depends on the relative advantage of the sperm/pollen compatibility allele on a given female/stylar genotype, 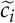 (that is, the relative fitness of the compatible sperm/pollen haplotype *M* on a the *i*^*th*^ female genotype), weighted by the frequency of the *i*^*th*^ diploid female genotype at the ℱ locus (i.e. before migration), *p*_*i*_ :

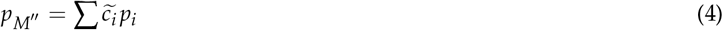

Where 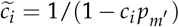. If the female (stylar) barrier is dominant, *c*_*FF*_ = *c*_*F f*_ = *c* and *c* _*f f*_ = 0, and 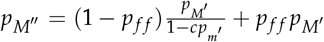. We can then find Δ_*p*_ fertilization as

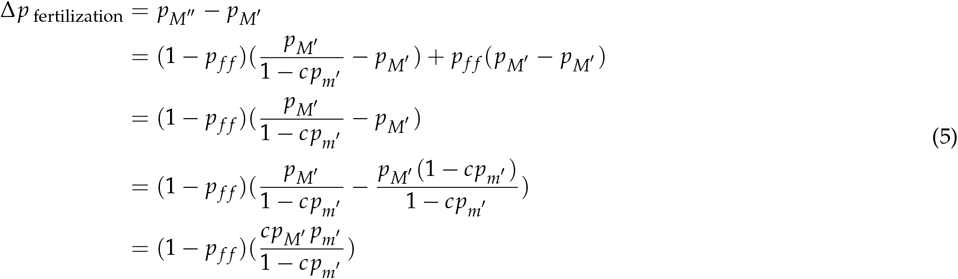

Equation 5 is the standard allele frequency change from population genetics weighted by the frequency with which selection can operate — i.e. the frequency of incompatible females.

#### Linked selection during fertilization

We now consider the change in frequency of an allele (*A* or *F*, described below as *X* which includes both alleles) in linkage disequilibrium with the sperm/pollen incompatibility allele *M* due to the fertilization advantage of *M*.

After mating, we calculate the frequency of paternally-derived haplotypes bearing this linked allele, *p*_*X*_ ″, where the double prime indicates that the frequency is calculated after both migration and mating. This quantity depends on the advantage of the sperm/pollen compatibility allele on a given female (stylar) genotype and the statistical association between alleles *M* and *X* (described by gametic phase linkage disequilibrium, *D*_*MX*_, which is the covariance between *M* and *X* alleles in sperm/pollen after sperm/pollen migration), weighted by the marginal frequency of the diploid genotype at the ℱ locus in females/styles (i.e. before migration), *p*_*i*_.

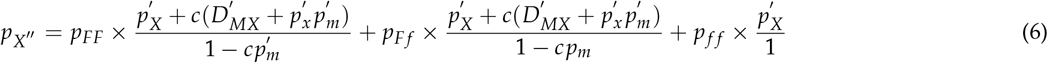

The change in frequency of an allele linked to ℳ between pollination (mating) and fertilization is therefore

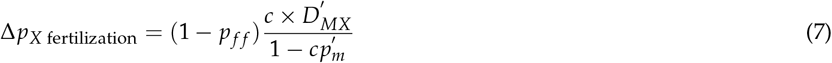

As such, the alleles in positive linkage disequilibrium with the sperm/pollen compatibility allele *M* always increase in frequency during fertility selection because of the fertilization advantage, so long as there are some incompatible females/styles in the population. Because linkage disequilibrium between *M* and *F*, as well as *M* and *A*, is never less than zero, all teosinte alleles increase in frequency in both populations by fertility selection (Figure 3B).

#### The generation of *trans* linkage disequilibrium during fertilization

We begin with two diverged populations, fixed for alternative alleles at local adaptation and sperm/pollen compatibility loci and with different allele frequencies at the female/stylar incompatibility. As such, LD between sperm/pollen compatibility and locally adaptive alleles is initially very large, and is broken down by migration and recombination.

This incompatibility system generates *trans* LD between sperm/pollen and female (stylar) loci — that is, compatibility rules generate a statistical association between paternally-derived *M* alleles and maternally derived *F* alleles. Before selection, this *trans* LD equals

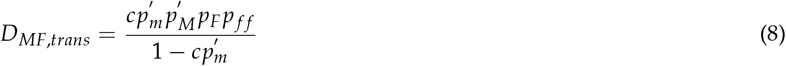

Clearly, this also indirectly generates *trans* LD between maternally-derived female incompatibilities and loci in LD with the sperm/pollen compatibility allele (or deviations from Hardy-Weinberg equilibrium at the *F* locus). *Trans* LD is converted to *cis* LD by recombination.

#### Local adaptation

Genotype frequencies at the local adaptation locus A after selection follow the standard population genetic equation 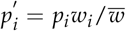

With multiplicative fitness effects, at a single local adaptation locus in the population in which *A* is favored, genotypic fitnesses are: *w*_*AA*_ = 1, *w*_*Aa*_ = (1 − *s*), and *w*_*aa*_ = (1 − *s*)^2^, and mean fitness equals 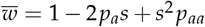. After selection, genotype frequencies are:

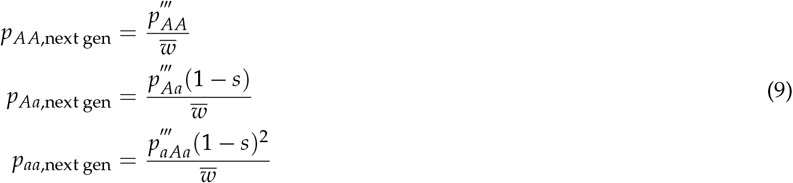

and the change in genotype frequencies between fertilization and viability selection are

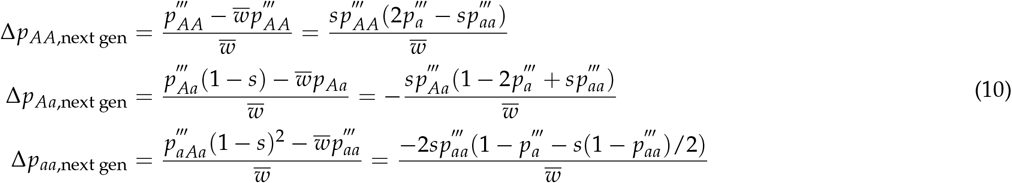

With multiplicative fitness, the locally maladapted allele always decreases by direct selection, regardless of allele frequency or inbreeding coefficient (Fig. S7).

#### Linked selection during local selection

After fertilization, selection directly changes the frequency of the local adaptation loci as described above. This also changes the frequency of loci statistically associated with the locally adapted allele. The change in frequency of an allele, *X* (e.g. *F* or *M*), linked to 𝒜 by linked selection is determined by the selection coefficient *s* and zygotic LD, and equals:

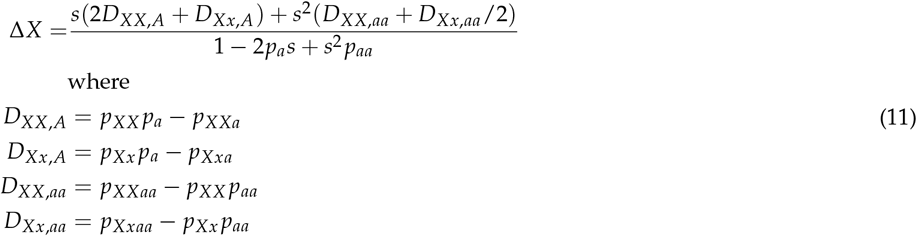

**Figure S1.**
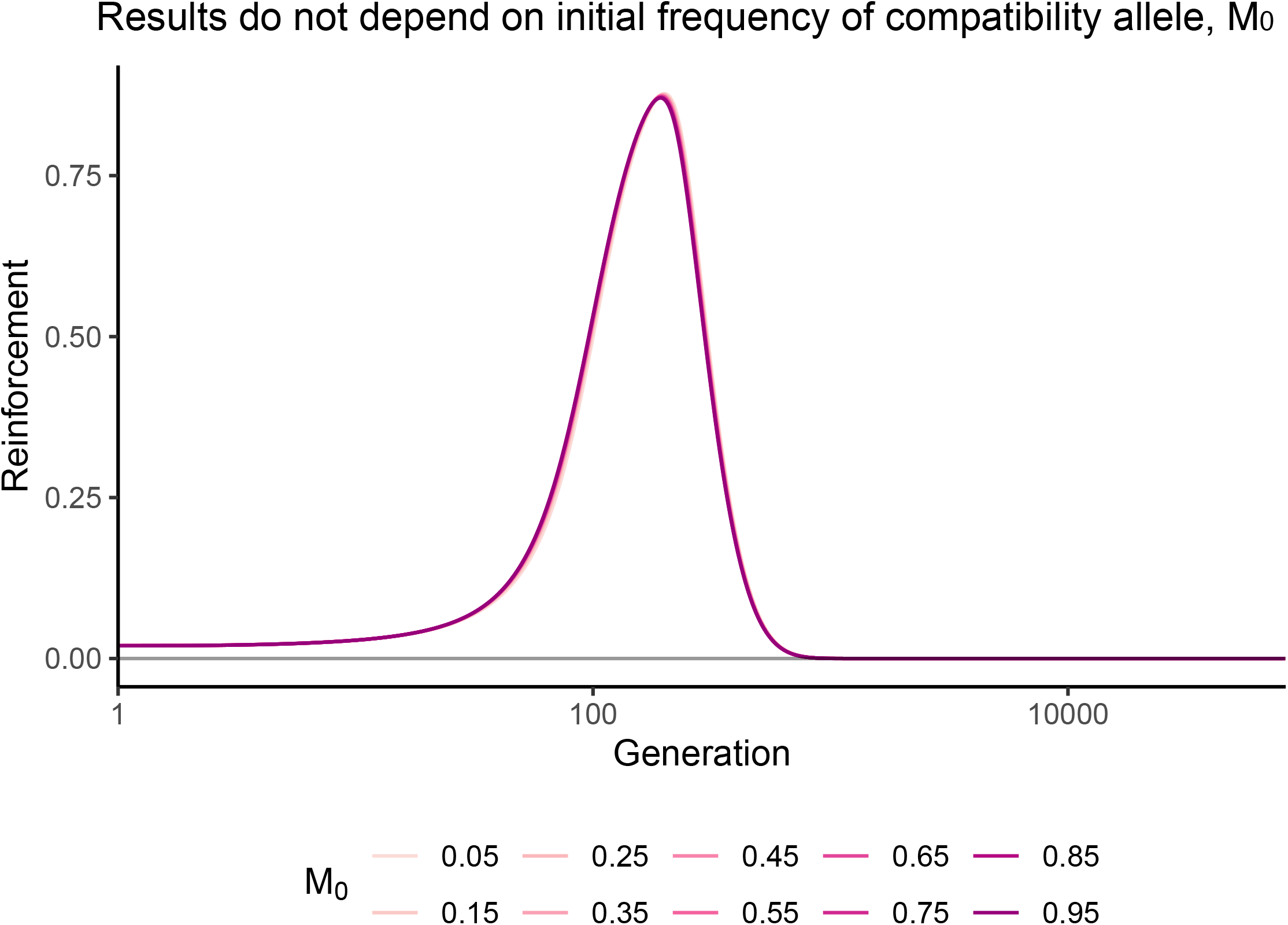
The initial frequency of *M* in the reinforcing species does not influence qualitative results. The impact of variability in the initial frequency of sperm/pollen compatibility allele, *M*, in the *reinf* on the transient reinforcement of postmating prezygotic isolation. All lines overlap. Parameter values: Selection — *s*_reinf_ = _*s*non-reinf_ = 0.75. Migration — (_*g*non-reinf→reinf_ = _*g*reinf→non-reinf_ = 0.1). Recombination — *r*_𝒜ℳ_ = *r*_ℳ ℱ_ = 0.0001. Allele frequencies — *f*_*M*0,*reinf*_ = displayed by color, *f*_*M*0,NR_ = 0, *f*_*F*0,R_ = 0.01, *f*_*F*0,NR_ =0.

**Figure S2.**
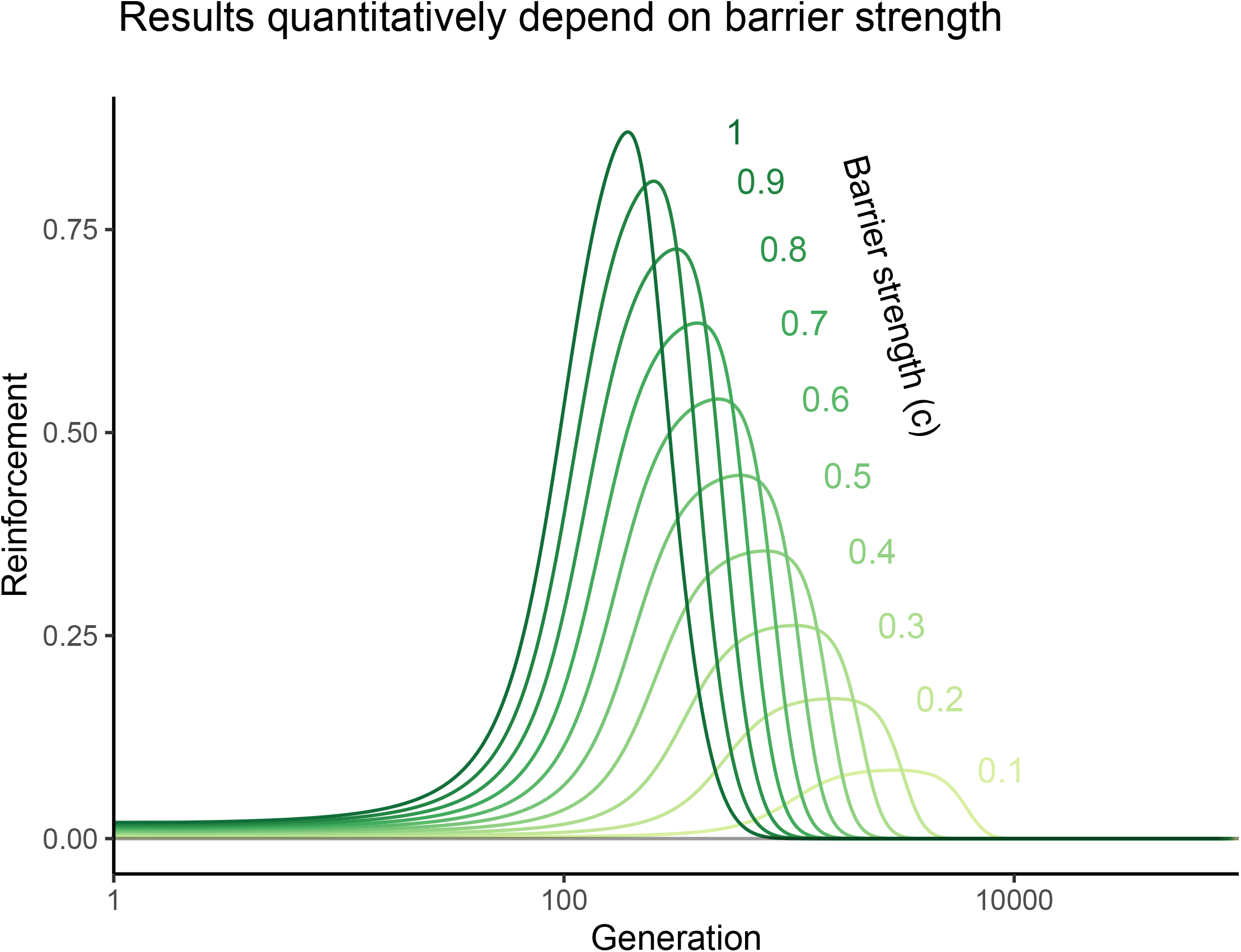
Female choosiness alters strength of reinforcement. We allow for an imperfect barrier (i.e. variation in female choice) by allowing females with fertilization barrier genotypes to be fertilized by a given haplotype, *k*, with probability 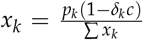, where *p*_*k*_ is the frequency of haplotype *k* in pollen after fertilization. *δ*_*k*_ equals zero for compatible sperm/pollen grains and one for incompatible sperm/pollen grains. *c*, the efficacy of the barrier, is colored in the plot above. Parameter values: Selection — *s*_R_ = *s*_NR_ = 0.75. Migration — (_*g*non-reinf→reinf_ = *g*_R→NR_ = 0.1). Recombination — *r* _𝒜ℳ=_ *r*_ℳ ℱ_ = 0.0001. Allele frequencies — *f*_*M*0,reinf_ = 1, *f*_*M*0,non-reinf_ = 0, *f*_*F*0,reinf_ = 0.01, *f*_*F*0,non-reinf_ = 0.

**Figure S3.**
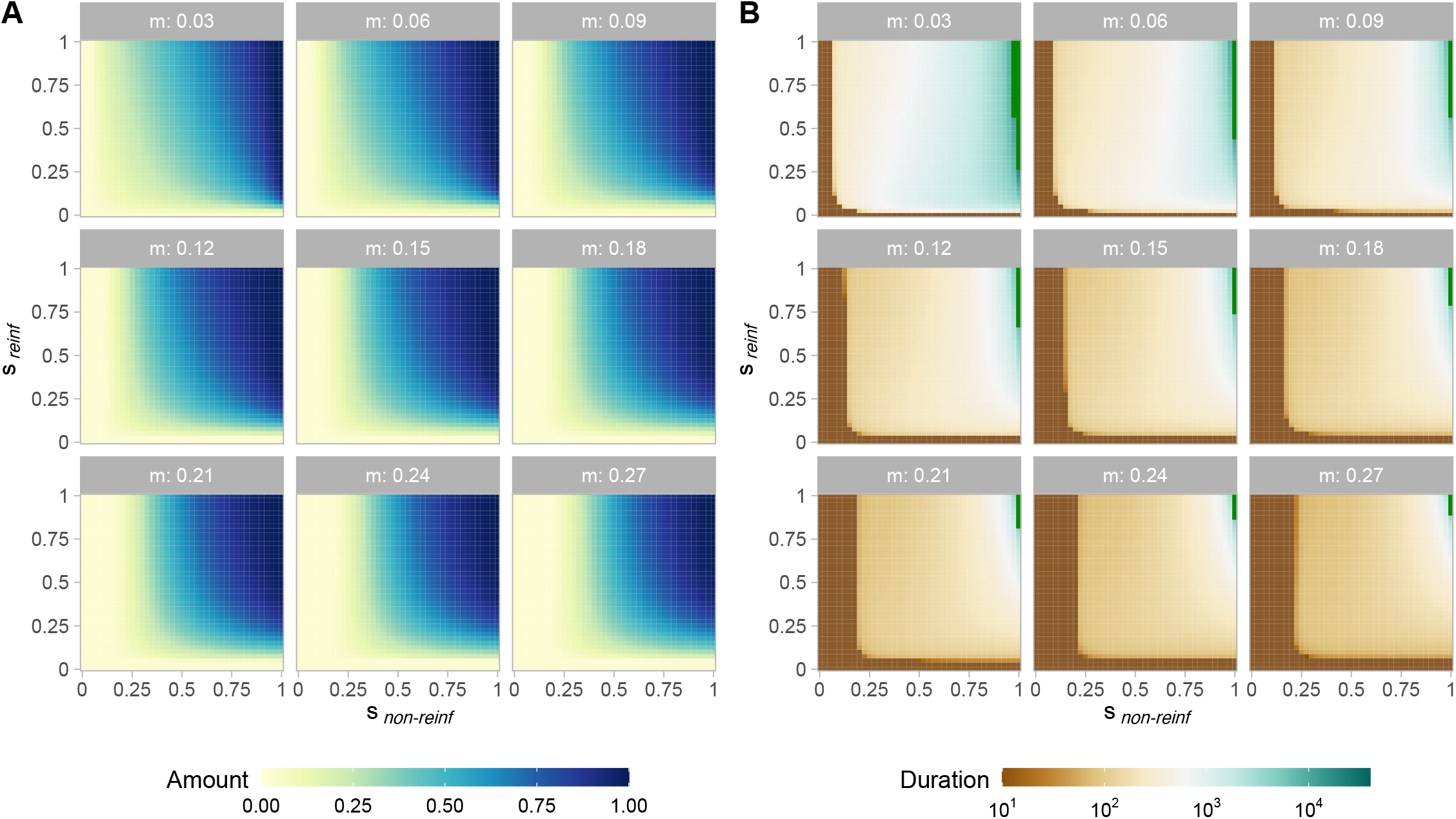
The impact of asymmetric selection on the extent (A) and duration (B) of reinforcement. Reinforcement strength and duration are estimated over a range of symmetric migration rates with *r*_𝒜ℳ_ = *r*_ℳ ℱ_= 10^− 4.^

**Figure S4.**
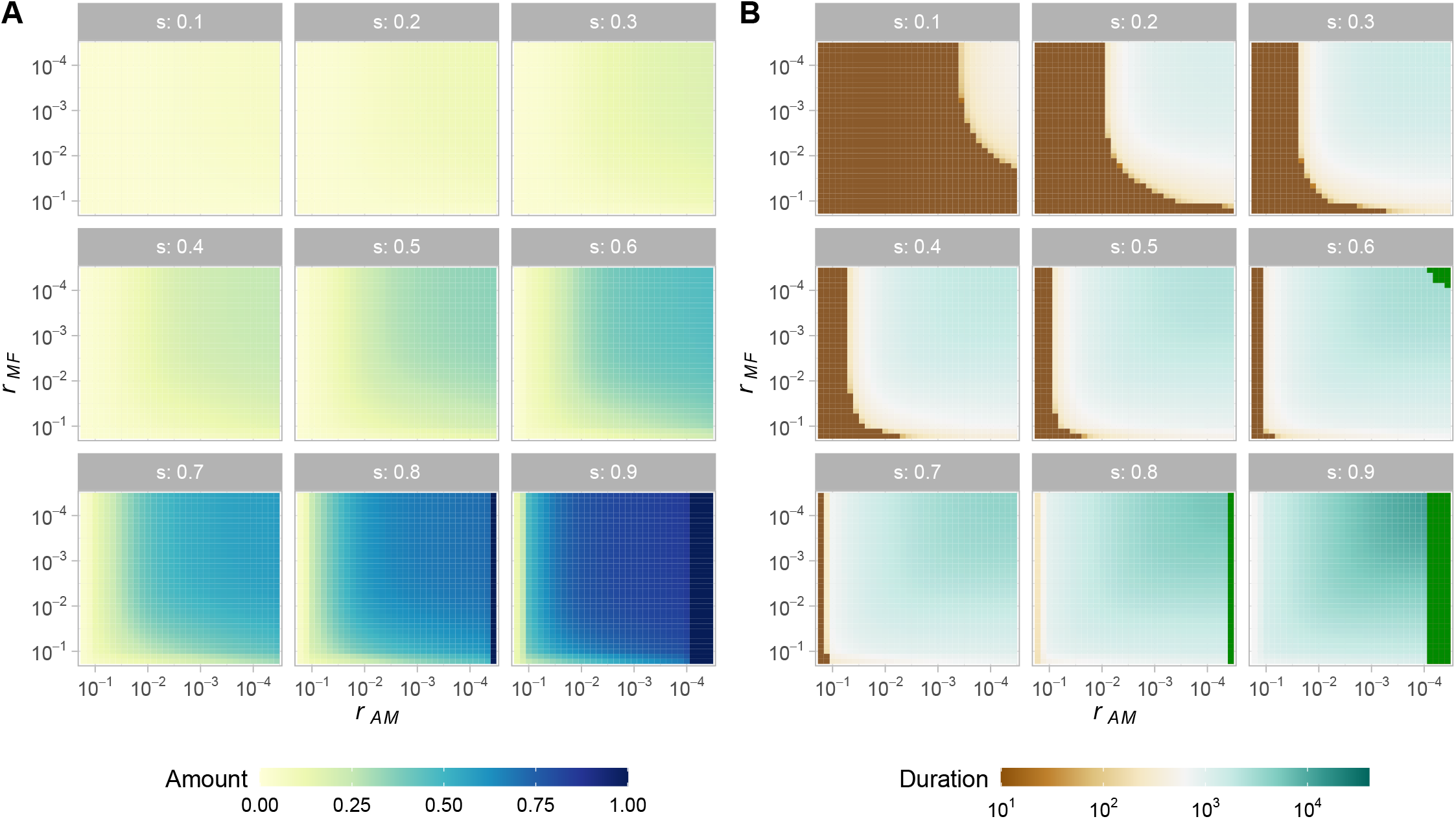
The impact of linkage on the extent (A) and duration (B) of reinforcement. Reinforcement strength and duration are estimated over a range of symmetric selection coefficients. *g*_non-reinf → reinf_ = *g*_reinf → non-reinf_ = 0.03. Note that the x-axis moves from loose linkage on the left to tight linkage on the right.

**Figure S5.**
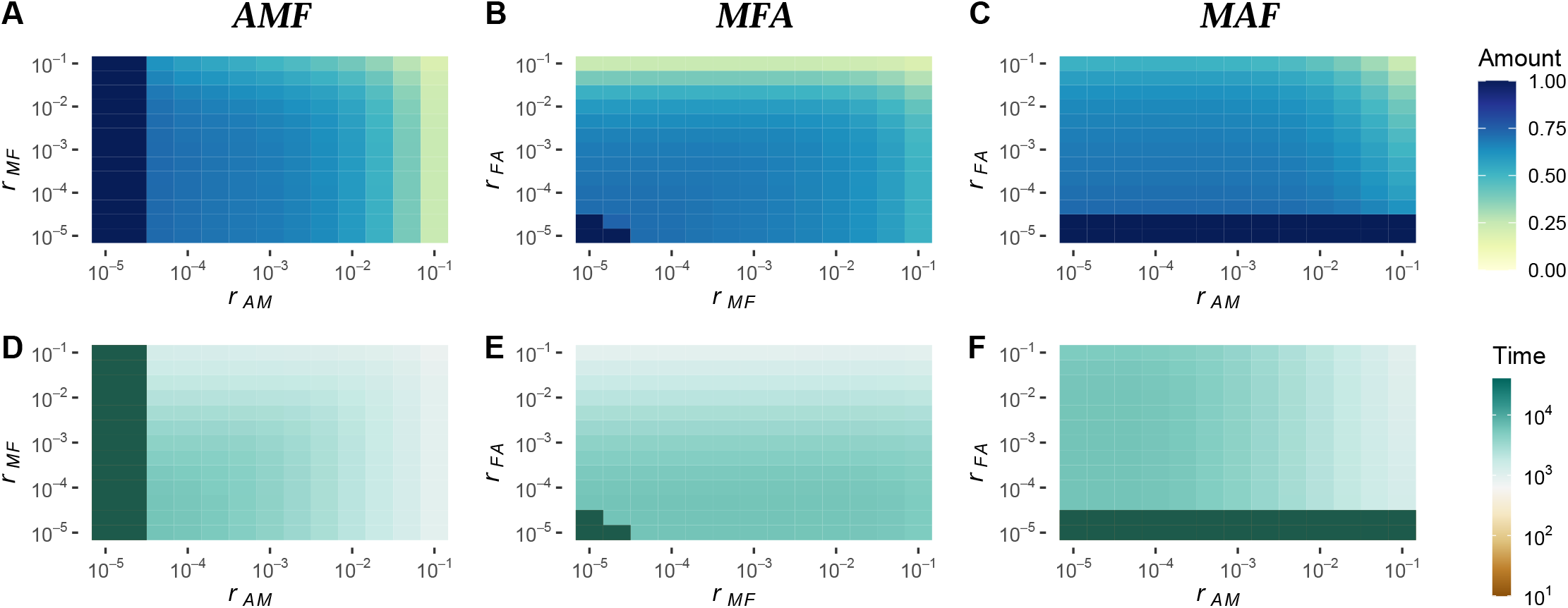
Locus order impacts the amount and duration of reinforcement. Top row (**A - C**) is reinforcement amount; bottom row (**D - F**) is duration, as estimated under different marker orders. Default marker order is 𝒜ℳ ℱ: amount (**A**); duration (**D**). Marker order MFA: amount (**B**); duration (**E**). Marker order ℳ 𝒜ℱ: amount (**C**); duration (**F**). Shown are results with a symmetric selection coefficient of 0.8 and migration *g*_non-reinf→reinf_ = *g*_reinf→non-reinf_ = 0.01.

**Figure S6.**
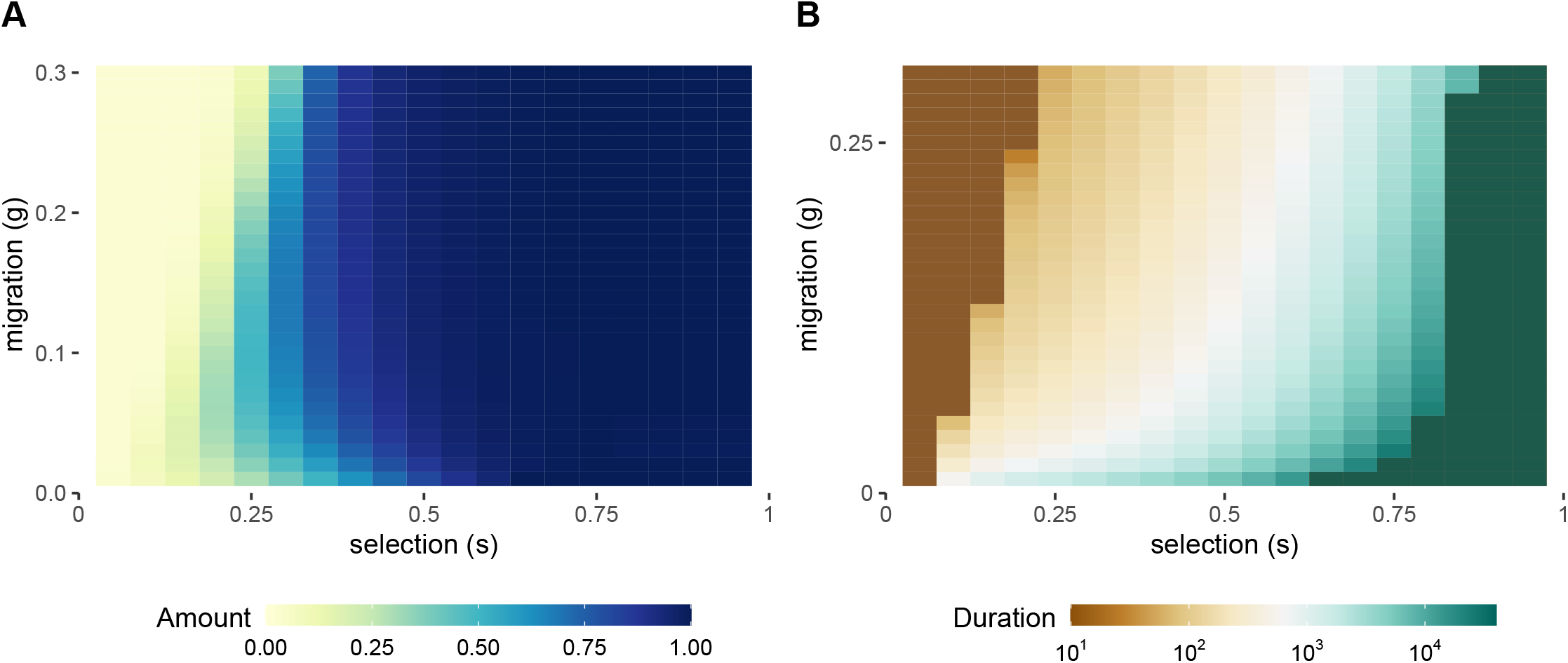
Asymmetrical variation in female preference does not underlie transience of reinforcement. Incorporating a second gametophytic factor in the non-reinforcing NR population does not qualitatively change the amount (**A**) or duration of reinforcement (**B**), although reinforcement begins at lower intensities of selection and reaches completion across more selection coefficients.

**Figure S7.**
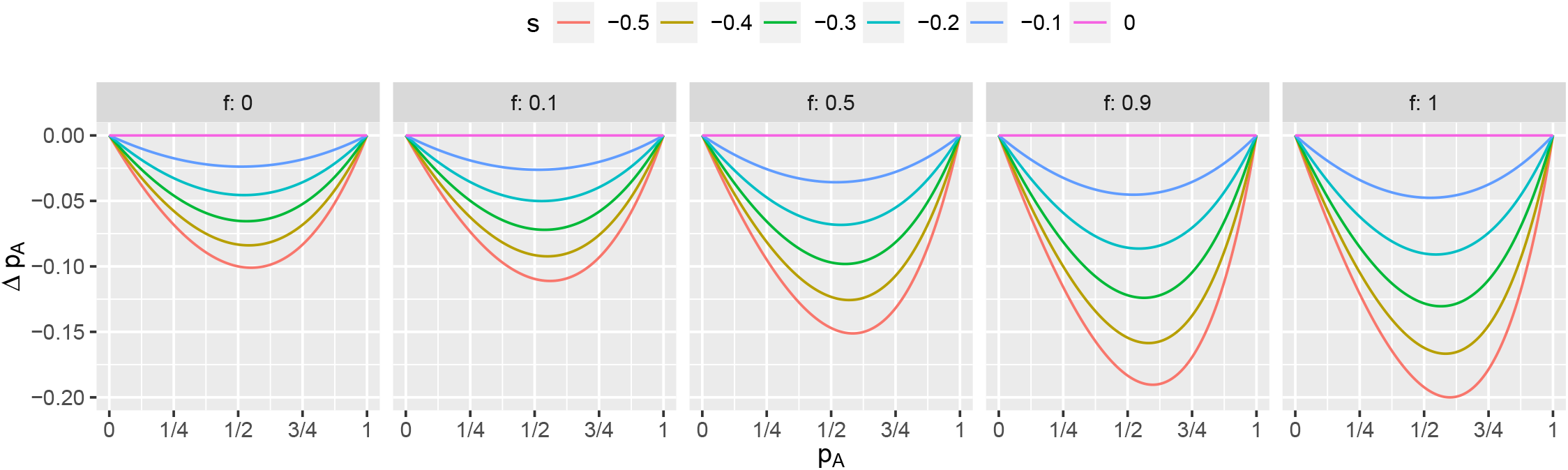
Allele frequency change at 𝒜 by direct selection. The locally maladaptive allele always decreases in frequency by direct selection *s*, regardless of the degree of assortative fertilization and/or population structure, summarized as the inbreeding coefficient *f*.

**Figure S8.**
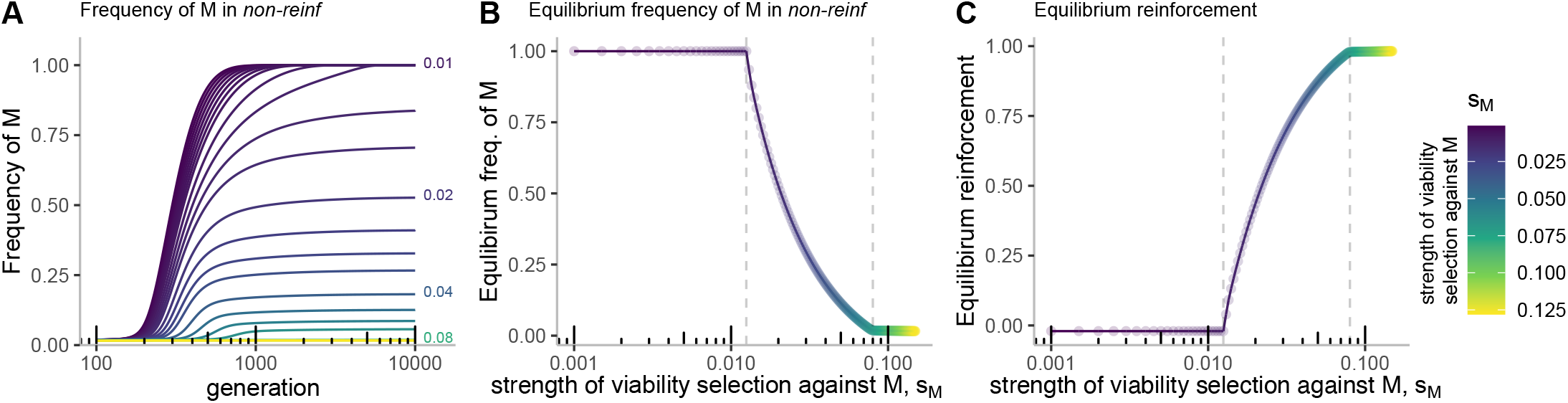
Introducing a cost (*s*_*M*_) to the male compatibility allele impacts the evolution of conflict over reinforcement. (**A**) The frequency of the *M* allele in the *non-reinf* over time as a function of the additive cost of the allele (*s*_*M*_ noted by color, with several values labeled for clarity). The frequency of the *M* allele in *non-reinf* (**B**) and the extent of reinforcement (**C**) after 10,000 generations (taken to be the equilibrium value, as evidenced by Figure S8A). Results are plotted on a *log*_10_ scale for clarity, though numbers are non-transformed. Dashed grey lines in (**B**) and (**C**) note the values (0.0125 and 0.08) at which we transition from transient reinforcement to a polymorphic equilibrium, and then to complete and stable reinforcement, respectively. All measures describe populations after selection and before recombination. This figure illustrates a single set of parameter values with one adaptive locus. Selection: *s*_reinf_ = *s*_non-reinf_ = 0.75; Migration: *g*_non-reinf→reinf_ = *g*_reinf→non-reinf_ = 0.1; Recombination: *r*_𝒜ℳ_ = *r*_ℳℱ_ = 0.0001; Allele frequencies: *f*_*M*0,R_ = 1, *f*_*M*0,non-reinf_ = 0, *f*_*F*0,reinf_ = 0.01, *f*_*F*0,non-reinf_ = 0. NR indicates the non-reinforcing population/species.

